# A strategy to study intrinsically mixed folded proteins: The structure in solution of ataxin-3

**DOI:** 10.1101/275131

**Authors:** A. Sicorello, G. Kelly, A. Oregioni, J. Nováček, V. Sklenář, A. Pastore

## Abstract

It has increasingly become clear over the last two decades that proteins can contain both globular domains and intrinsically unfolded regions which both can contribute to function. While equally interesting, the disordered regions are difficult to study because they usually do not crystallize unless bound to partners and are not easily amenable to cryo-electron microscopy studies. Nuclear magnetic resonance spectroscopy remains the best technique to capture the structural features of intrinsically mixed folded proteins and describe their dynamics. These studies rely on the successful assignment of the spectrum, task not easy per se given the limited spread of the resonances of the disordered residues. Here, we describe assignment of the spectrum of ataxin-3, the protein responsible for the neurodegenerative Machado-Joseph disease. We used a 42 kDa construct containing a globular N-terminal josephin domain and a C-terminal tail which comprises thirteen polyglutamine repeats within a low-complexity region. We developed a strategy which allowed us to achieve 87% assignment of the spectrum. We show that the C-terminal tail is flexible with extended helical regions and interacts only marginally with the rest of the protein. We could also, for the first time, deduce the structure of the polyglutamine repeats within the context of the full-length protein and show that it has a strong helical propensity stabilized by the preceding region.

## Introduction

It is now well established that proteins can be classified into three large families. Some proteins adopt a well-defined stable tertiary structure and are usually referred to as globular proteins. Others are devoid of a three-dimensional structure but can adopt it when and if needed. This family is described as that of intrinsically unfolded proteins (IUPs or IDPs). A third family is composed of proteins containing both globular and intrinsically unfolded regions. These proteins could be considered as ‘mixed folded proteins’ (MFPs). The study of all three families is important and offers specific challenges to researchers. Of the three main methodologies able to describe structures at atomic resolution, X-ray crystallography, nuclear magnetic resonance (NMR) and more recently cryo-electron microscopy, NMR has proven most helpful in the characterization of IDPs and MFPs, because it can operate directly in solution, capture molecular motions and observe flexible structures. There are however several difficulties which disseminate these studies. The first and possibly most important one is due to the fact that, when not helped by the chemical shift dispersion of the NMR spectrum of a folded protein, IDP regions suffer of an intrinsic and inevitably inherent severe overlap of the NMR resonances. Since the assignment of an NMR spectrum to each of the atoms along the sequence depends on the tracing of the magnetization transfer, that is the transfer from one atom to the next as in a network, overlap has the consequence of limiting the possibility of uniquely tracing the chain. Several strategies have been suggested to circumvent this problem (1-3). Yet, no definite answer has been given to the question and new ideas must be considered to eventually find, if possible, a unique and universal strategy.

Here, we describe how we have gained significant insights into the structure of ataxin-3, a human 42 kDa protein involved in the neurodegenerative disease spinocerebellar ataxia type-3 or Machado Joseph disease (4). Ataxin-3 is a typical example of an MFP: it contains an N-terminal 21 kDa domain, josephin, followed by a highly flexible region containing two or three (depending on the isoform) ubiquitin interacting motifs (UIMs) and a tract of polymorphic polyglutamine (polyQ) repeats (5). The disease is thought to be caused by aggregation of the protein promoted both by the josephin domain and by the polyQ tract (6, 7). The age of onset and the severity of the disease depends on the length of the polyQ tract and the disease reveals when this is above 42 repeats. Ataxin-3 is a deubiquitinating enzyme which recognizes and cleaves preferentially polyubiquitin (polyUb) chains of at least four subunits and is supposed to play an important role in the ubiquitin-proteasome pathway (8, 9). Ataxin-3 binds to ubiquitin through different surfaces, distributed both along the josephin domain and in the C-terminus (10,11).

Solving the structure of ataxin-3 by crystallography has proven difficult presumably because of difficulties in obtaining ordered crystals of the full-length protein. We resorted to NMR as the most promising strategy. However, we hit against considerable difficulties not only because of the long predicted unstructured regions which result in very low dispersion of the spectrum but also because of the presence of several low complexity stretches along the sequence. Additionally, ataxin-3 is prone to aggregate and this makes it difficult to keep the protein in solution for the period (days or weeks) required for the formation of crystals or for the acquisition of complex NMR datasets. Accordingly, sparse information both from X-ray and NMR has been published for specific regions of the protein but a comprehensive picture of ataxin-3 structure is still elusive.

Despite the intrinsic difficulties, we successfully developed a strategy which has allowed us to obtain assignment of 87% of the spectrum and gain a clear map of the ataxin-3 structure. We could clearly observe the secondary structure propensity of the unfolded and flexible chain. Within this, the polyQ tract has a strong tendency to a helical conformation stabilised by the preceding region. Our results provide the essential prerequisite for any further study of the interactions between ataxin-3 and other cellular partners and suggest new tools for the study of MFPs.

## Materials and methods

### Construct choice and cloning

The plasmid used for ataxin-3(Q13) recombinant over-expression is a derivative of pMAL-c5x (NEB), hereafter referred as pMht-atx3(Q13). The plasmid encodes the maltose binding protein (MBP) followed by a hexa-histidine tag, a tobacco etch virus protease (TEVp) cleavage site and the sequence encoding ataxin-3(Q13). The expression plasmid for the isolated josephin domain (pMht-jos) was obtained by replacing the codon encoding residue 183 of ataxin-3 with a TAA stop codon via inverse PCR site-directed mutagenesis. The mutants of ataxin-3(Q13) were obtained via inverse PCR site-directed mutagenesis using pMht-atx3(Q13) as a template. The desired mutation was inserted at the 5’-end of the forward mutagenic primer. After PCR amplification the linearised plasmid was purified using Zyppy™ Plasmid Miniprep Kit (Zymo Research). The DNA was then incubated with DpnI (New England Biolabs), T4 polynucleotide kinase (New England Biolabs) and QuickLigase (New England Biolabs) to degrade the template, phosphorylate the 5’ ends and re-circularise the plasmid, respectively. The ligation mix was used to transform directly DH5α cells (New England Biolabs) and the cells were plated overnight on LB/agar plates at 37°C. The outcome of the mutagenesis was checked by Sanger sequencing (GATC Biotech).

### Protein production

The isolated josephin domain, ataxin-3(Q13) wild type and mutants were produced in a recombinant form using the pMht series plasmids described in the previous section and the *E*. *coli* strain BL21. Slightly different protocols were needed for producing the uniformly or selectively labelled samples. ^15^N-^13^C and ^15^N uniformly labelled proteins were expressed in M9 medium containing ^15^N ammonium sulphate and ^13^C-D-glucose as the sole source of nitrogen and carbon respectively. The cells were grown at 37°C until an OD_600_ of 0.8 was reached. Protein over-expression was induced by adding isopropyl β-D-1-thiogalactopyranoside (IPTG) to a final concentration of 1 mM. The cells were further incubated under shaking at 37°C for 3 h.

^15^N-amino acid selective labelling was achieved by growing bacterial cells in M9 medium supplemented with ^14^N ammonium sulphate, ^14^C glucose and 0.1 g of each unlabelled amino acid. When the cells reached an OD of 0.8-1.0, 100 mg of the desired ^15^N labelled amino acid and 1 g of all other unlabelled amino acids were added to 1 litre of culture. The cells were incubated for 15-20 min under shaking. For ^15^N-glutamine selective labelling, 75 mg of 6-Diazo-5-oxo-L-norleucine, 180 mg of L-methionine sulfoximine, 180 mg of L-methionine sulfone were also added to the culture. For ^15^N-glutamate selective labelling, 1 g of L-methionine sulfoximine, 1 g of L-methionine sulfone, 250 mg of disodium succinate, 250 mg of disodium maleate, 250 mg of aminooxyacetate were added to the culture. After incubating the cells under shaking at 37°C for 15-20 min, IPTG was added to a final concentration of 1 mM and protein over-expression was allowed for 90 min.

All cultures were harvested by centrifugation at 6000*xg* for 20 min on a Beckman Avanti centrifuge. The cell pellet was transferred to 50 ml tubes and frozen at −20°C. The cell pellet was defrosted and resuspended in 20 mM sodium phosphate, 200 mM NaCl, pH 7.5 (buffer A) supplemented with DNAse and lysozyme. After sonication, ataxin-3 was purified by NiNTA-agarose (Generon) affinity chromatography using 20 mM sodium phosphate, 200 mM NaCl, 250 mM imidazole, pH 7.5 (buffer B) for elution. The 6xHis-MBP tagged protein was then buffer exchanged to buffer A and hexa-histidine tagged Tobacco etch virus protease was added to release untagged ataxin-3(Q13). The cleavage mixture was incubated with NiNTA-agarose resin to remove 6xHis-MBP and 6xHis-TEVp. The flow-through containing ataxin-3(Q13) was concentrated with centrifugal filters and run on a Superdex 75 26/60 size exclusion chromatography (SEC) column (GE Healthcare) to remove small traces of aggregates or impurities and exchange the protein into 20 mM sodium phosphate buffer, 2 mM TCEP, pH 6.5. The elution volumes obtained from SEC confirmed that all the proteins were monomeric in solution. A final purity higher than 95% was assessed by SDS-PAGE.

### NMR spectroscopy and sequential assignment

All NMR experiments were performed in 20 mM sodium phosphate buffer at pH 6.5 and 2 mM TCEP at 25°C. Data were acquired at 600 MHz, 700 MHz, 800 MHz and 950 MHz on Bruker Avance instruments equipped with cryogenic probes. The HNCO, HN(CA)CONH and HabCabCONH experiments were acquired with four scans per increment. The HNCO was acquired with an interscan delay of 1.5 s, spectral widths of 10416.7 Hz (acquisition dimension) 2594.7 Hz (^15^N) 1810.94 Hz (^13^C’), The non-uniform sampling of the indirectly detected dimensions was applied. The 5D HNcaCONH experiment was acquired with an interscan delay of 1.5 s, spectral widths of 10416.7 Hz (acquisition dimension), 2594.8 Hz (^15^N), 1811.1 Hz (^13^C’), 2594.8 Hz (^15^N) and 960.2 Hz (^1^H), The maximal evolution times were 50 ms for t1, t2 and t4 (^1^H and ^15^N) and 26.5 ms for t3 (^13^C’). The 5D HabCabCONH experiment was acquired with an interscan delay of 1.5 s and with spectral widths of 10416.7 Hz (acquisition dimension), 2594.8 Hz (^15^N), 1810.9 Hz (^13^C’), 12474.8 Hz (^13^C_aliph_) and 6401.2 (^1^H_aliph_). The maximal evolution times were 10 ms for the ^1^H_aliph_, 6.7 ms for ^13^C_aliph_, 26.4 ms for ^13^C’, and 49.3 ms for ^15^N indirect dimensions. In both experiments, a total number of 1910 points was acquired in the indirect dimensions. A whole dataset of HNCO, HNcaCONH and HabCabCONH experiments was acquired on the same protein sample in ∼65 h. Additional ^15^N-^1^H HSQC experiments were recorded immediately before and after each higher dimensionality experiment to assess the spectral integrity of the protein. In all cases, the two HSQC spectra did not show discrepancies suggestiong sample stability which is necessary to acquire high quality data of high dimenstionality.

HSQC based pseudo-3D relaxation data were acquired at 800 MHz. For the isolated josephin domain, delays of 100, 200, 400, 700, 1200, 1800, 2500 ms and 16, 32, 64, 96, 144, 192 ms were used for T_1_ and T_2_ measurements, respectively. For ataxin-3(Q13) delays of 100, 200, 400, 700, 1000, 1500, 2500, 4000 ms and 16, 32, 48, 64, 96, 128, 160, 192, 240 ms were used for T_1_ and T_2_ measurements, respectively. Average T_1_ and T_2_ values for josephin and for ataxin-3(Q13) were calculated by fitting the area under the spectra in the range 7.7-10.5 ppm to an exponential function. The relaxation rates for josephin residues within ataxin-3(Q13) were obtained by fitting the HSQC peak intensities. The correlation times were estimated using the formula 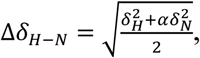.

The spectra were processed with NMRPipe based scripts and NMRDraw (12). The HNCO and the 5D experiments were processed with the Multidimensional Fourier Transform (MFT) (13,14) and the Sparse Multidimensional Fourier Transform (SMFT) (15) algorithms. The processing software was downloaded from the authors’ website. The assignment of the NMR spectra was performed with the software Sparky 3.115 (T.D. Goddard and D. G. Kneller, University of California, San Francisco, USA) and CcpNmr Analysis 2.4 (16). The relaxation rates were calculated and analysed using nmrglue (17), nmrPipe and CcpNmr Analysis 2.4. The assignment was submitted to the BMRB database (accession number 27380).

## Results

### The spectrum of full-length ataxin-3

Assignment of full-length ataxin-3 was performed on isoform 2 (Uniprot ID: P54252-2) with an interrupted tract of 13 polyQ repeats, Q_3_KQ_10_-hereafter referred to as ataxin-3(Q13). This isoform contains three UIMs in the C-terminus at variance with isoform 1, used in previous studies of our group (5,18), which has only two UIMs. Two features of ataxin-3 had to be considered when choosing a suitable assignment strategy. Since ataxin-3 tends to aggregate with time, ^15^N-^1^H HSQC spectra were recorded before and after each experiment to assess whether the protein remained mainly in a monomeric state throughout the experiment. The spectra were assessed for variations of the peak volumes and of the resonance positions. Typically, a fresh sample of ataxin-3(Q13) could be used for at least three days in the NMR tube at 25°C. No significant changes in both peak intensity and position could be detected over this time-frame. The sequence of the C-terminus of ataxin-3(Q13) (residues 183-361) contains multiple repetitive stretches including three UIMs and the polyQ tract. The HSQC spectrum of ataxin-3(Q13) clearly shows the coexistence of well dispersed resonances with regions characterised by poor dispersion and high resonance overlap (**Figure 1**). Spectral overlap and crowding were encompassing the frequency range 117-126 ppm (in the ^15^N dimension) and 7.9-8.7 ppm (in the ^1^H dimension). This is a hallmark of the intrinsic high flexibility of the C-terminus as expected from the sequence. Another important feature is that the intensities of the resonances are highly uneven, with many resonances being extremely broad. This suggests different dynamical properties.

**Figure 1.**
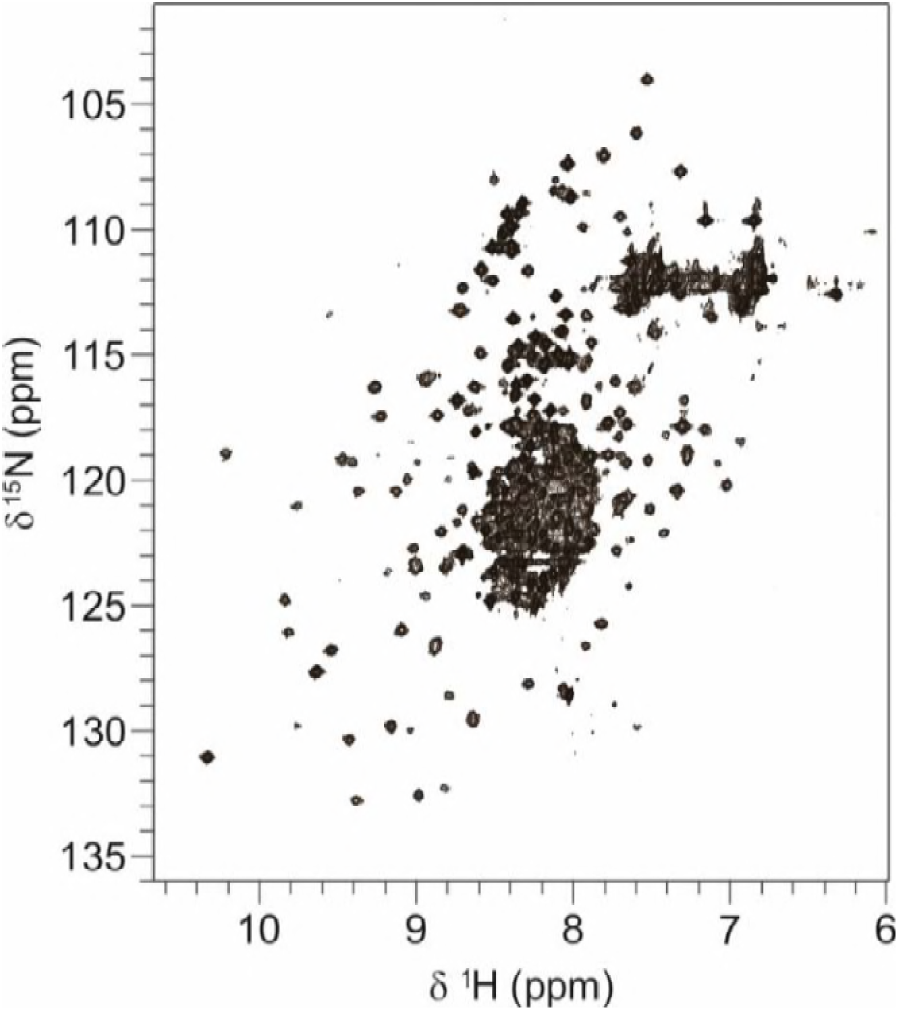
^15^N-^1^H HSQC spectrum of ataxin-3(Q13). The spectrum was recorded at 800 MHz and 25°C.

Taken together, these data show that the tail of ataxin-3 has all features of an intrinsically unfolded or partially structured system.

### The josephin domain does not significantly interact with the C-terminus

To identify the already assigned resonances of the smaller josephin domain (residues 1-182) within the much more crowded spectrum of ataxin-3, we superimposed the ^15^N-^1^H HSQC spectra of isolated josephin with that of full-length ataxin-3(Q13) (**Figure 2A**). Most of the well dispersed josephin resonances in the spectrum of full-length ataxin-3 overlap with those of the isolated domain. This allowed facile identification of several resonances. Some peaks, however, shift due to a change in the chemical environment. This could be a consequence of either interaction between the structured and unstructured regions or of minor changes of the experimental conditions. While the latter effect can be minimal in a well dispersed protein spectrum, unambiguous assignment of a spectrum of this complexity becomes prohibitive without further support. Initially, we could tentatively assign 124 out of the 174 non-proline resonances of josephin (70%). Each of the N-H correlations in the spectrum of the isolated josephin domain was used to guide the search for the corresponding resonance of the full-length protein. The identity of the N-H resonances of josephin in ataxin-3(Q13) was further confirmed by inspection of an HNCACB spectrum. Given the unfavourable tumbling time of the josephin domain in the context of the full-length protein, this experiment yielded weak signals for the Cα and Cβ and no correlations with the preceding residue for most of josephin NH resonances. This partial information proved nevertheless very valuable in circumscribing the amino acid type, yielding unambiguous assignment in most cases.

**Figure 2.**
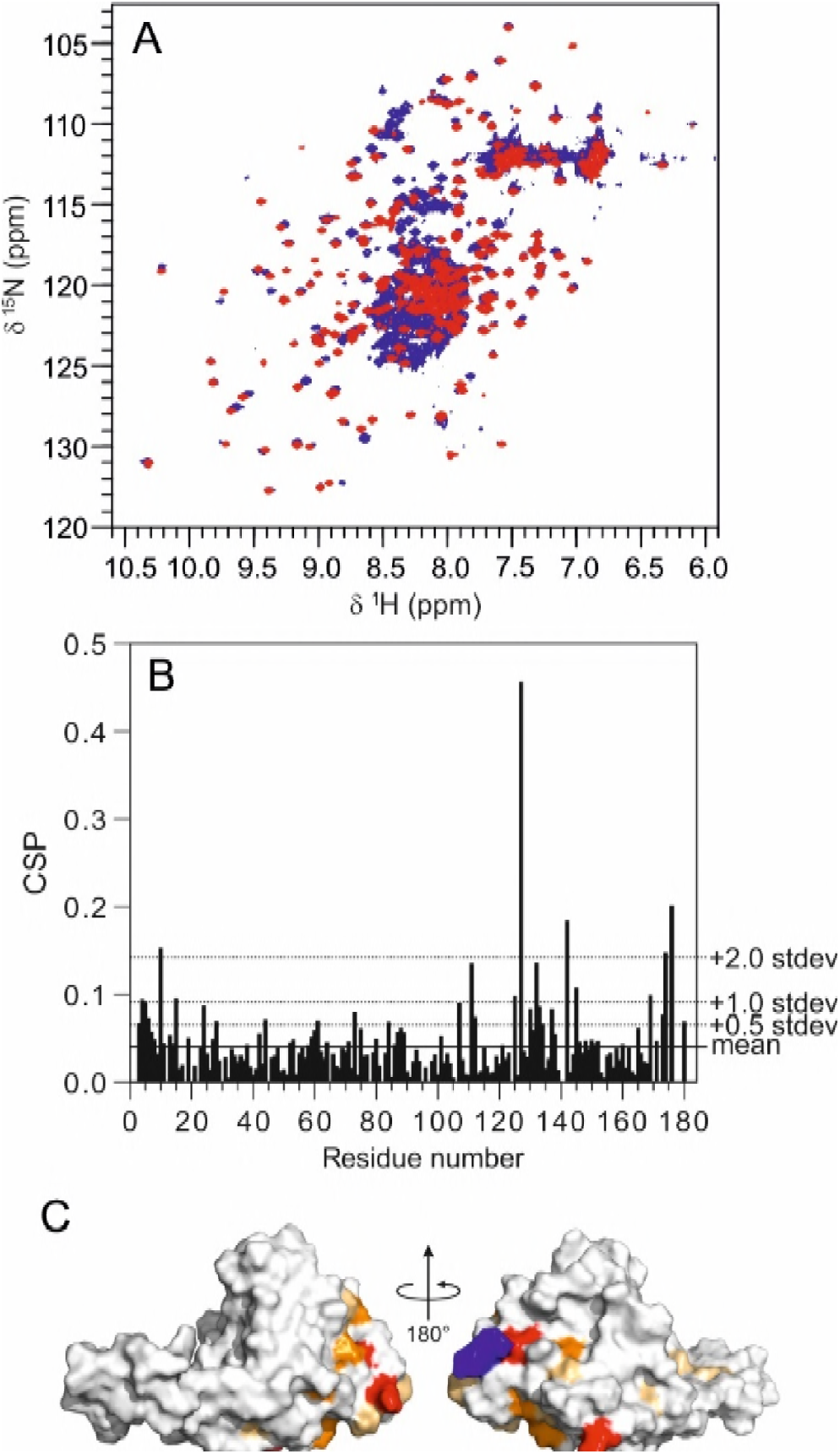
Assignment of the NH backbone resonances of the josephin domain within the ataxin-3(Q13) construct. A) Superimposition of the ^15^N-^1^H HSQC spectrum of the isolated josephin domain (red) and ataxin-3(Q13) (blue). B) Chemical shift perturbation of josephin NH backbone resonances in ataxin-3(Q13) with respect to the isolated domain. The chemical shift perturbations (Δδ_H-N_) were weighted using the formula 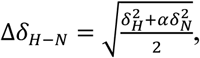 where α = 0.14 for glycine and α = 0.20 for any other residue. C: mapping of the CSPs on the structure of the josephin domain. Light orange: CSP > average + 0.5 standard deviations; Orange: CSP > average + 1 standard deviations; Red: CSP > average + 2 standard deviations. The C-terminal arginine is coloured in blue.

To extend further the assignment of the josephin resonances within the full-length ataxin-3(Q13) spectrum, we compared the 3D NOESY-HSQC spectra of ataxin-3(Q13) and isolated josephin domain (**Fig. S1 in the Supporting Material**). This allowed assignment of 20 additional josephin resonances, reaching a sequence coverage of 83% of the josephin resonances. A few josephin resonances in the HSQC spectrum of ataxin-3(Q13) showed very low signal-to-noise ratio and did not yield carbon correlations in the HNCACB spectrum or NOESY-HSQC amide or aliphatic proton patterns which could be identified in the spectrum of the isolated josephin domain. To identify these resonances, we acquired a BEST-TROSY HSQC (19) and overlapped it with the conventional HSQC spectra of ataxin-3(Q13) and the isolated josephin domain. The overall increase in signal-to-noise of the BEST-TROSY HSQC spectrum made it possible to detect the previously invisible N-H josephin resonances and assign them. The combination of these experiments helped us to obtain an assignment coverage of 91% of the josephin spectrum which corresponds to a ∼55% coverage of full-length ataxin-3. These results strongly indicate the importance for the sequential assignment of difficult proteins of exploiting a range of different experiments designed to fulfil different purposes. This includes experiments which would not usually be considered as a first choice for protein assignment, such as the NOESY-HSQC.

The chemical shift perturbations observed in josephin within ataxin-3(Q13) as compared to the isolated domain are small (overall within a standard deviation below or equal to 0.5) (**Figure 2B**). The only resonances with more appreciable variations are in the N-and C-termini and at residue G127. This is reasonable because the N-and C-termini of josephin are close in space. G127 is spatially close. This evidence demonstrates convincingly lack of tight interactions between the josephin domain and the C-terminal tail of ataxin-3.

### Walking through the assignment of ataxin-3 C-terminus

Triple resonance experiments (HNCA, HNCACB, CBCAcoNH) were initially performed to assign the ataxin-3(Q13) C-terminus (**Figure S2**). The low dispersion of the ^1^H/^15^N/^13^C resonances associated to residues within repetitive regions impeded progress in the assignment, suggesting the necessity of ultra-high resolution techniques. We exploited a strategy developed for intrinsically disordered proteins which uses 5D pulse sequences (20). The high resolution resulting from the higher dimensionality of these experiments has been shown to be particularly successful for systems that are not only intrinsically unfolded but also highly degenerate in sequence (21-23). In this strategy, a HN(CA)CONH experiment is used to provide sequential connectivity by linking the ^15^N-^1^H chemical shifts of consecutive residues, while a HabCabCONH spectrum is used to identify amino acid types from their Cα and Cβ resonances (20). The data are acquired using a non-uniform sampling (NUS) scheme in which only 2% of the total data-points are collected. The spectra are subsequently reconstructed using Sparse Multidimensional Fourier Transform (15). This strategy resulted in a reduction of the total acquisition time needed for a full set of assignment experiments from weeks to ∼2.5 days, a time frame in which the ataxin-3 spectral properties are unperturbed, indicating that the protein remains mainly monomeric. We first acquired a reference high resolution NUS HNCO spectrum which was used to extract the chemical shift values for each ^1^H/^15^N/^13^C’ resonance from the parent CON and CabHab 2D planes of the 5D HNcaCONH and HabCabCONH datasets respectively. The CON plane contains a weak resonance related to the HNCO peak of residue *i* and a stronger resonance related to the HNCO peak of residue *i-1*. The CabHab planes contain the Cα, Cβ, Hα and Hβ resonances of residue *i-1*. The 5D spectra were of excellent quality and confirmed the initial assignment obtained via triple resonance experiments. Additional residues could be assigned, resulting in the coverage of ∼50% of the resonances of ataxin-3 C-terminus (**Figure 3**). Assignment of the ^1^H, ^15^N and ^13^C’ resonances was possible for residues 214-226, 240-246, 263-274, 298-312, 319-360. Within these regions, the Cα chemical shifts were assigned for all the residues with the exception of residues 214, 218, 220, 227, 246, 268, 273, 274, 306, 310, 312, 324, 327, 331, 339, 359, and 360, for which it was not possible to detect a C’ resonance in the sequential assignment. The Cβ chemical shifts could be assigned for all the residues with the exception of residues 227, 246, 274, 312, and 360 for which no signals could be detected in the CabHab planes of the 5D HabCabCONH experiment.

**Figure 3.**
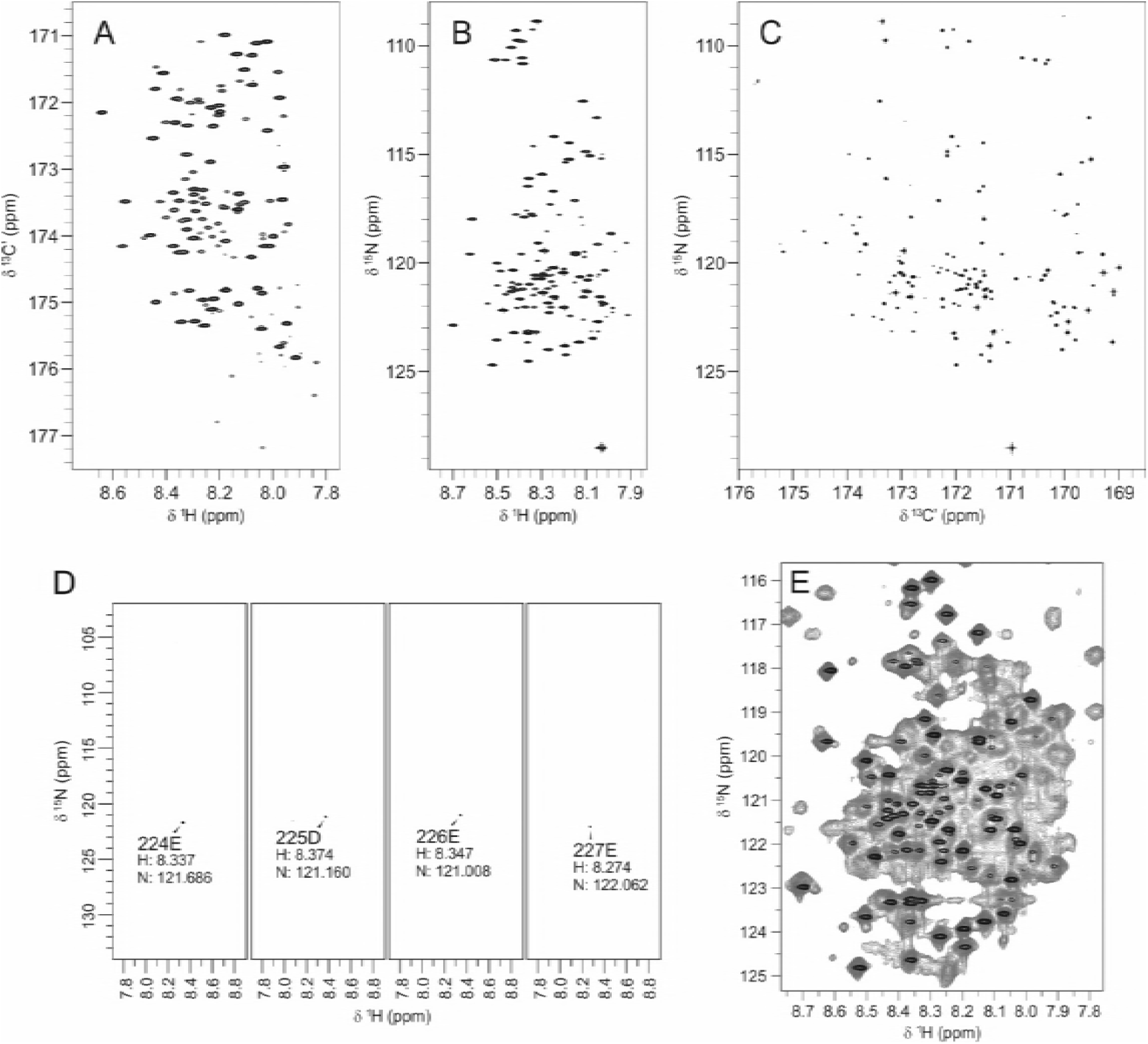
Assisting assignment with 5D HNcaCONH experiments. A) ^13^C-^1^H, B) ^15^N-^1^H and C) ^15^N-^13^C projections of the HNCO experiment. D) ^15^N-^1^H planes of consecutive residues associated to HNCO peaks in the 5D HNcaCONH experiment. E) Superimposition of the HSQC spectrum of ataxin-3(Q13) (grey) with the ^15^N-^1^H projection of the HNCO spectrum used as reference for the 5D HNcaCONH (black).

In the course of this analysis, we noticed that only a subset of the expected resonances could be detected. 5D experiments are optimized for highly disordered systems. The long polarization transfer pathway used in these experiments allows ultra-high resolution but also results in a considerable loss of signal-to-noise. Consequently, fast relaxing resonances located in rigid or semi-rigid portions of the polypeptide chain are usually non-detectable, while the effect is much less pronounced for the slowly relaxing resonances from highly flexible regions. Therefore, the mere fact that only a subset of the expected resonances could be detected in the 5D experiments indicates that the residues in the C-terminal tail of ataxin-3 have a wide range of dynamical properties. In particular, the ultra-high resolution of high dimensionality experiments allowed sequential assignment of 8 out of 13 glutamine residues in the polyQ tract (residues 298-305). The signal-to-noise of the carbon resonances in this region progressively increases from the N-terminus to the C-terminus. This suggests a higher flexibility of the polyQ tract at the C-terminus simply based on this observation.

### Systematic mutation of key residues

Additional resonances could be assigned by integrating the information obtained from the 5D experiments and from triple resonance experiments. Despite this, the information was still insufficient to obtain a satisfactory assignment coverage. For some amide resonances, a correlation with the previous or following residue was impeded by the absence of the corresponding Cα_i-1_ in the HNCACB spectra. An additional problem was posed by the ambiguity of many Cα-Cα_i-1_ correlations, leading to multiple equally likely candidates for sequential assignment. To overcome these problems, a systematic mutation strategy was exploited. Mutations were introduced at different sites within both assigned and unassigned regions. HSQC spectra were acquired for each mutant that could be purified and was stable. This assignment strategy relies on the fact that the resonance in the spectrum of the wild-type protein that is associated to the mutated residue will not be detectable in the spectrum of the mutant, while a resonance with different ^1^H and ^15^N chemical shifts, associated with the new residue, is expected to appear. Since the C-terminal tail of ataxin-3 is flexible and does not seem to form stable tertiary structures, a mutation at one site of the protein is expected to cause a chemical shift perturbation only for the amide NH resonances of residues proximal to the mutation. It is reasonable to expect that this perturbation is roughly inversely correlated with the distance in sequence from the mutated site in a flexible and largely solvent exposed chain. The mutants G193A, T207A, L213I, D228E, S256A, S260A, R282H, R284H, R285H, and C316A were successfully used to assign resonances for which a through-bond correlation could not be detected in triple resonance experiments. For example, the analysis of the HSQC spectrum of the mutant T207A in conjunction with the triple resonance experiments of the wild-type protein allowed unambiguous assignment of residues 206-208, based on the combination of the chemical shift perturbation observed and on information about the amino acid type from Cα and Cβ resonances obtained from the HNCACB experiment.

### Further assignment using amino acid selective labelling

An amino acid selective labelling approach was used to resolve the ambiguities caused by the overlap of resonances associated with residues whose β carbons share similar chemical shifts.^14^N-^1^H labelled ataxin-3(Q13) samples were produced which incorporated a single ^15^N-labelled amino acid (^15^N-alanine, ^15^N-isoleucine, ^15^N-leucine, ^15^N-lysine, ^15^N-methionine, ^15^N-valine, ^15^N-arginine, ^15^N-glutamine, ^15^N-glutamate, ^15^N-tyrosine and ^15^N-phenylalanine). The samples resulted in an overall high labelling yield and selectivity, suggesting that amino acid metabolic scrambling was suppressed efficiently. Only the spectrum of selectively labelled ^15^N glutamate showed considerable scrambling from glutamate to glutamine, aspartate and alanine and some isotope dilution. Despite this, a critical analysis of the spectrum helped by comparison with the spectra of the other amino acid selectively labelled spectra helped discerning selective labelling from scrambling. This allowed unambiguous identification of amino acid types throughout the spectra (**Table S1**).

The assignment of overlapping resonances for which the chemical shifts of Cα_i-i-1_ and Cβ_i-i-1_ connectivity could be detected in HNCACB spectra was straightforward. When the Cα_i-1_ and Cβ_i-1_ were not detectable, assignment was obtained by producing amino acid selectively labelled mutants. For each labelling, the HSQC spectrum of wild-type ataxin-3 was compared with the HSQC of the corresponding point mutant. In this way, we could identify unambiguously V183, H187, L191, E194-A197, Q202-V204, K206, E210, R231, A232, R237, Q238, R251, A252, Q254, L255, M257, Q258, R262, E279-F289, and R318 (**Figure 4**).

**Figure 4.**
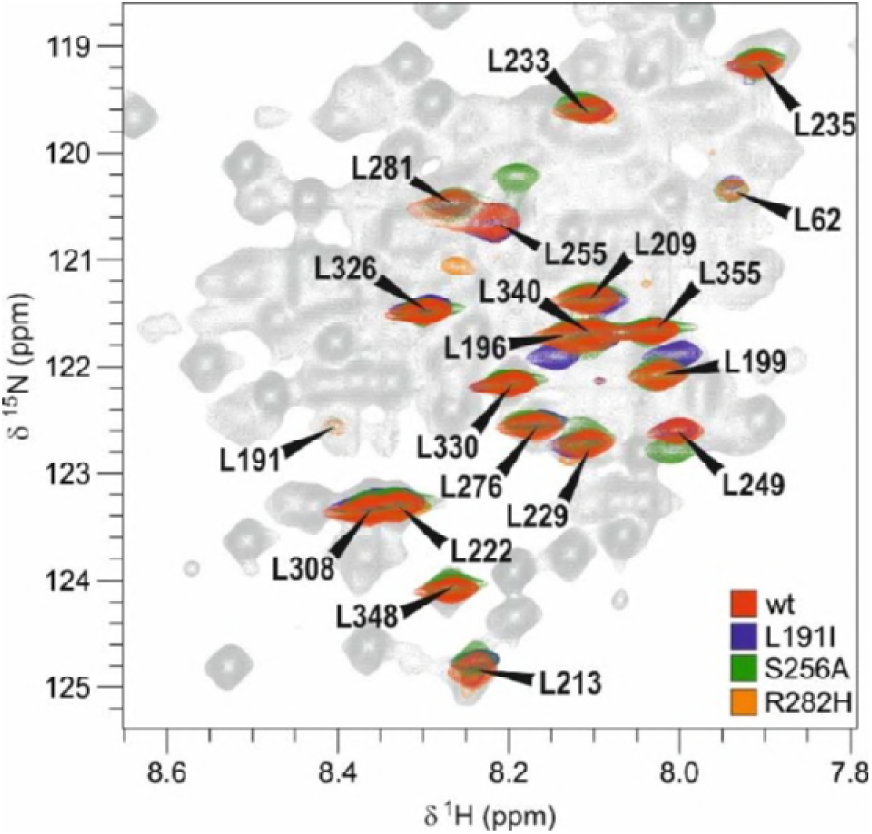
^15^N-^1^H HSQC spectra of ^15^N-Leu selectively labelled wild-type and mutated ataxin-3(Q13). The spectrum of the wild-type protein (red) is superimposed to the spectra of three mutants (L191I: blue, S256A: green, R282H: orange). The counter levels were adjusted to show only the sharper resonances related mainly to residues in the C-terminal tail. The resonances of at least eight more leucines were also detectable in the experiment but with lower intensities. They all correspond to Josephin leucines. Minor scrambling was observed at the noise level.

The NH resonances from most of the residues in the tract 183-192 could not be detected in the spectra of selective amino acid labelled samples. This fact and the lack of NH resonances of residues in the spatially close N and C-termini of the josephin domain suggest that this region is in an intermediate exchange regime.

### The use of BEST-TROSY HNCACB to assist assignment

In an attempt to increase the signal-to-noise further, band-selective excitation short-transient (BEST) experiments were performed in association with transverse relaxation-optimized spectroscopy (TROSY) using the BEST-TROSY HNCACB pulse sequence (24). This experiment achieves a longitudinal relaxation optimization allowing fast pulsing and rapid acquisition of spectra with higher resolution and signal-to-noise. The combination of BEST with TROSY enhances further these effects. It has been shown that the BEST sequence enhances preferentially resonances from transiently-structured regions which have usually a low signal-to-noise. The increased signal-to-noise of the BEST-TROSY-HNCACB experiment was used to confirm weak carbon resonances throughout the sequence of ataxin-3. This technique allowed, for instance, to detect the Cα^i-1^ and Cβ^i-1^ correlations of residues E279 and E280 to the preceding serine and glutamate, respectively. The combination of this information with glutamate selective labelling of wild-type ataxin-3 and of the R282H mutant led to unambiguous assignment of these residues.

Almost all the detectable resonances in the HSQC spectrum could be assigned unambiguously, reaching a coverage of the C-terminus of ataxin-3(Q13) of 158 out of 176 (90%) non-proline residues and the 87% of the full-length protein (**Figures 5,6**).

**Figure 5.**
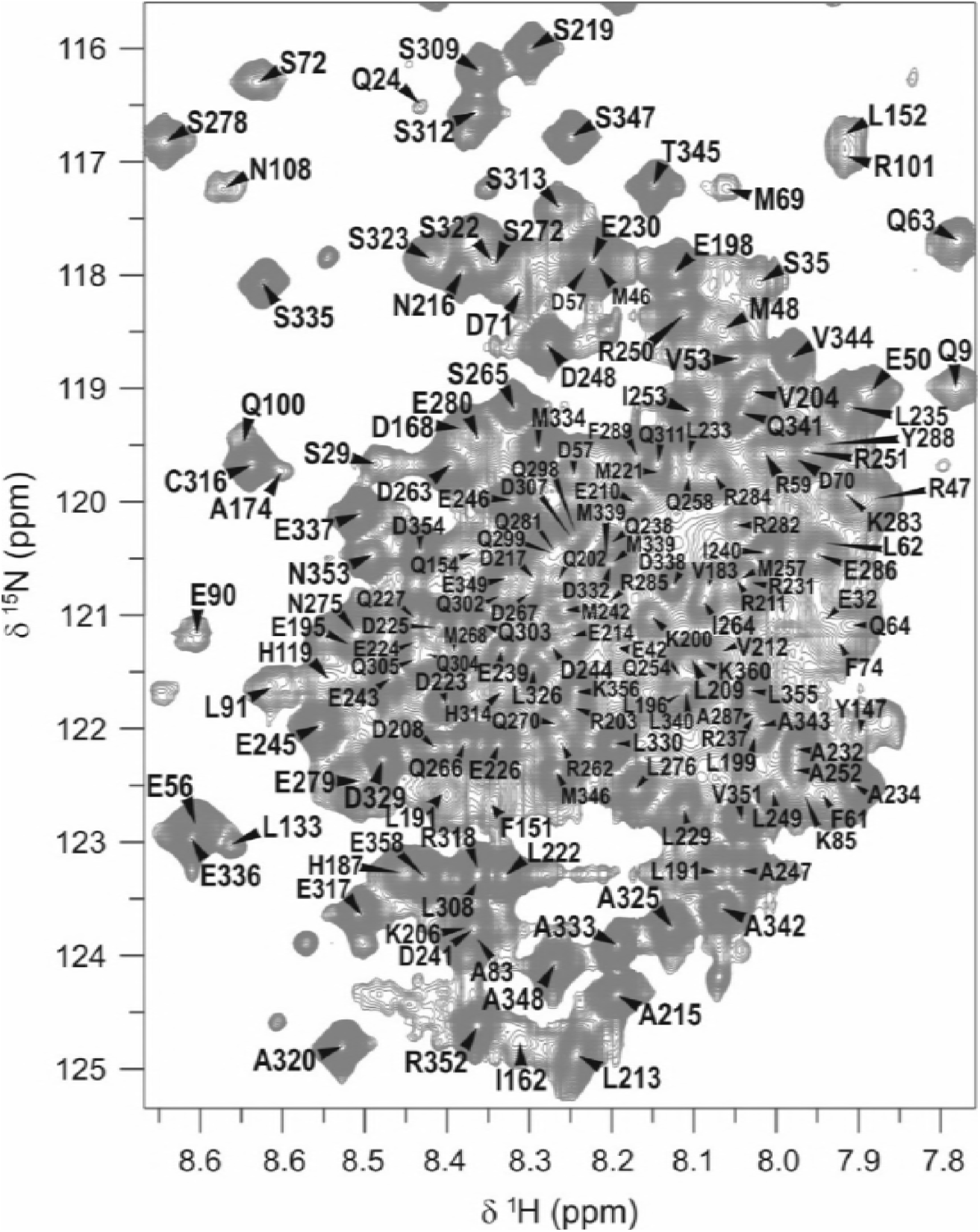
Assignment of the backbone NH resonances of the ^1^H-^15^N HSQC spectrum of ataxin-3 in the central, highly degenerate area of the spectrum.

**Figure 6.**
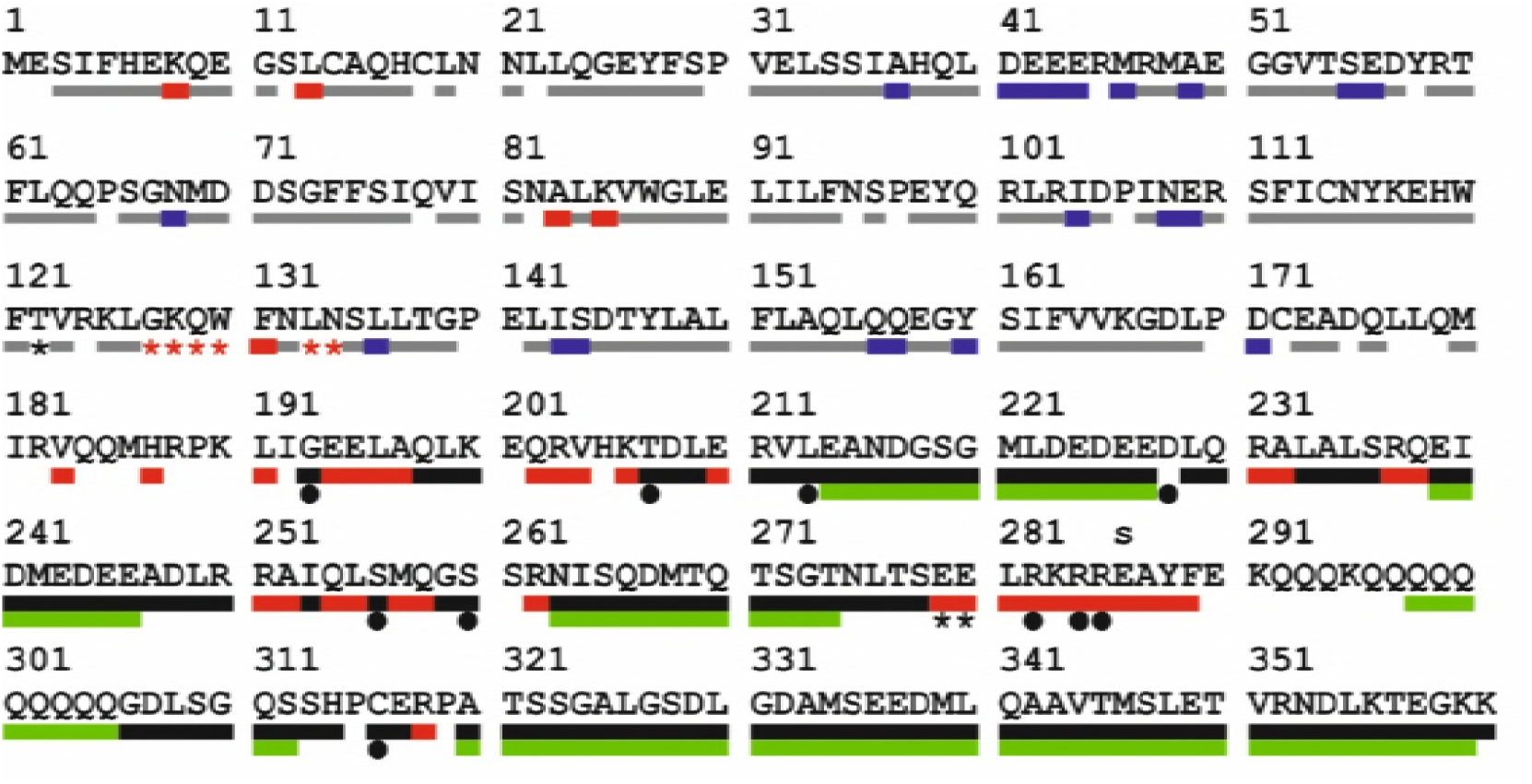
Assignment of the sequence of ataxin-3(Q13). The colour of the bars is associated to the technique used to aid the assignment. The backbone H pairs of the josephin domain (residues 1-182) were assigned by comparison with the assignment of the spectrum of the isolated domain without (grey bars) or with the aid of either ^15^N-amino acid selective labelling (red bars) or BEST-TROSY HSQC (red stars). Residues assigned by comparison of the NOESY-HSQC of ataxin-3(Q13) with that of the isolated josephin domain are indicated by blue bars. The C-terminal tail was assigned via HNCACB and CBCAcoNH (black bars), 5D HNcaCONH and HabCabCONH (green bars), selective ^15^N-amino acid labelling (red bars), BEST-TROSY-HNCACB (black stars) and single point mutations (black circles).

### Structure and dynamics of the C-terminal tail of ataxin-3

The Cα and Cβ chemical shits of the C-terminal tail of ataxin-3 allowed us to gain important information about the structure propensity and dynamics of this region which could not easily be obtained with any other method (**Figure 7A**). The algorithm SSP (25) generates a score indicating the propensity of each residue to populate helical, extended or random coil secondary structure motifs. Overall, the C-terminal tail is mostly highly flexible but with a strong helical propensity throughout with maximal values in the three UIMs (residues 224-240, 244-263, 331-348) in agreement with reports on peptides spanning the isolated UIMs (11). Interestingly, also the residues in the polyQ tract that could be assigned (298-305) exhibit some helical propensity. This is in support to the structure of a complex of a polyQ peptide with an antibody (26) but disagrees with our previous data showing that polyQ repeats C-terminally fused to a carrier protein (glutathione-S-transferase) are random coils independently of their length (27). Since however residues 278-289, which precede the polyQ repeats, have a helical propensity even higher than that of the UIMs, we can reasonably assume that the polyQ helicity is induced and stabilised by the preceding region. In agreement with this hypothesis is the fact that the α-helicity of the polyQ tract progressively decreases moving from Q298 to Q305.

**Figure 7.**
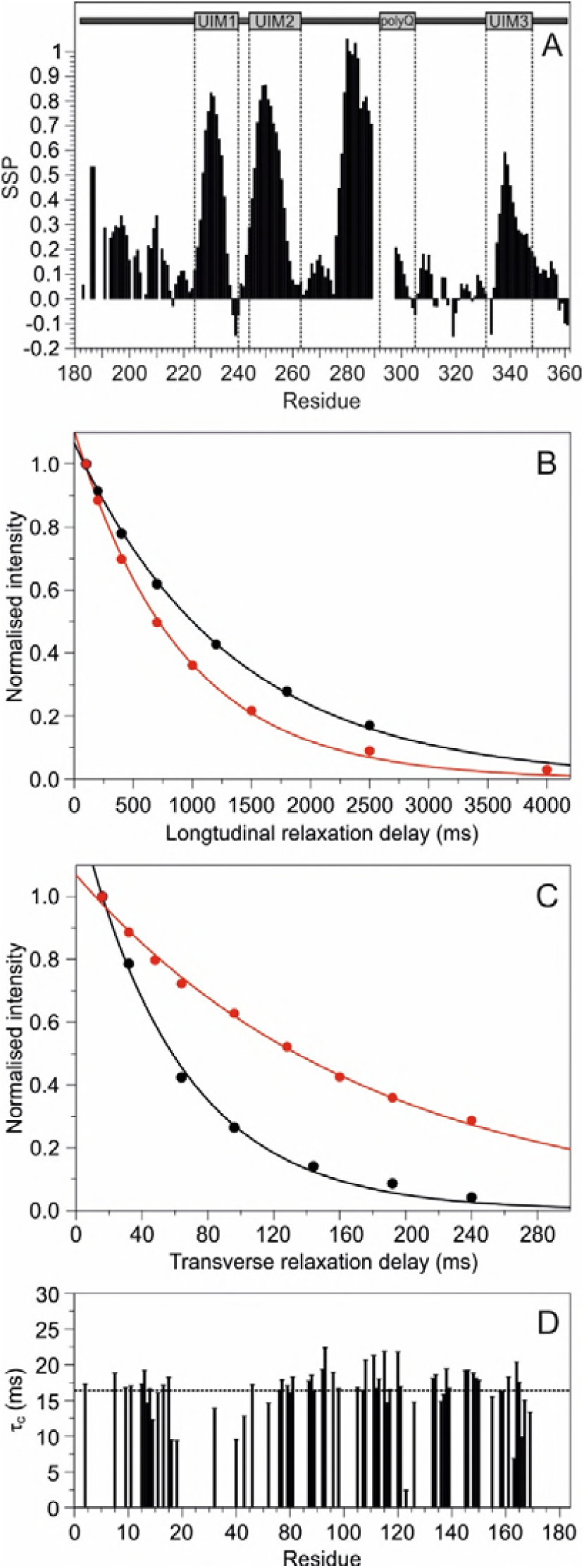
Description of the secondary structure and dynamics of ataxin-3(Q13). A) Secondary structure of the C-terminus of ataxin-3(Q13). SSP values around zero indicate random coil regions; positive and negative values indicate α-helical or β-sheet conformations respectively. B) Longitudinal and C) transverse relaxation of the spectrum of the isolated josephin domain (black) and ataxin-3(Q13) (red) in the range 7.7-10.5 ppm. D) Correlation times (τ_C_) of well dispersed resonances of josephin within the spectrum of ataxin-3(Q13). The dashed line indicates the average τ_c_.

The intrinsic flexibility of the C-terminal tail and dynamical properties very different from those of josephin are reflected also in the ^15^N-^1^H relaxation values (**Figure 7B,C,D**). The T_1_ and T_2_ relaxation times of the C-terminal tail, as estimated by fitting the area of the resonance envelop over the ^1^H region 7.7-10.5 ppm of the ataxin-3(Q13) spectrum, are 902.2 ms and 176.1 ms respectively, as opposed to values of 1322.8 ms and 61.3 ms for the isolated josephin domain (**Figure 7B,C**). Assuming a spherical and rigid polypeptide chain, the averaged correlation time obtained from the T_1_/T_2_ ratios of the isolated josephin domain is 10.9 ns. This value is in excellent agreement with the value previously reported (28) and corresponds to that expected for a protein with a molecular weight of ∼18 kDa (29). Under the same assumptions, we estimated an average value of 4.8 ns for full-length ataxin-3(Q13). This marked underestimation clearly indicates that the assumptions do not pertain this case and that the C-terminal tail is flexible and contributes by significantly lowering the apparent correlation time. To estimate the correlation time of josephin within ataxin-3(Q13), we selected a set of well-dispersed isolated resonances of josephin and measured their relaxation rates by traditional experiments (**Figure 7D**). The resulting averaged correlation time (16.4 ns) is that expected for a protein with a molecular weight of ∼25 kDa (29).

## Discussion

Ataxin-3 is an interesting protein not only because it is associated to the neurodegenerative Machado-Joseph disease but even more because it is a deubiquitinating enzyme with very specific fold and features (28, 30, 31). The first biophysical studies on this protein date around 2000 although much of the original interest was devoted towards the aggregation properties of the protein because these seem to be directly related to disease (6, 32, 33). In 2003, we proved that ataxin-3 is a MFP which contains an N-terminal josephin domain present in a restricted number of other eukaryotic species and a C-terminal tail (5). The first NMR structure of josephin was published in 2005 and validated the year after (28,34). From then on, however, no direct information became available for the ataxin-3 C-terminal tail. It is only now that we can report full assignment of the NMR spectrum of ataxin-3 thanks to a formidable assignment exercise.

Spectral assignment was indeed not trivial because of the several problems imposed by a MFP of this size: the NMR spectrum of ataxin-3 fully reflects a mixture of globular and unstructured regions. As such the assignment strategy was far from standard and we had to develop an *ad hoc* procedure which combined different independent routes. We found the application of 5D techniques indispensable to achieve the resolution necessary to assign highly degenerate regions. Despite this, the use of high dimensionality techniques was limited by the high dynamic heterogeneity and the presence of resonances in or near an intermediate exchange regime, as supported by selective labelling. Judicious introduction of site specific mutations helped us to push the assignment coverage and confirming the information obtained independently. Given the high complexity of the system, we considered this validation essential to exclude incorrect tracing.

Our effort resulted in an overall 87% coverage assignment which extends over the whole protein. While we cannot exclude that the unassigned residues have resonances perversely overlapping, independent evidence suggests the presence of conformational and/or solvent exchange of some residues especially in the region between josephin and the C-terminal tail. It was also paradoxically difficult to identify some of the josephin resonances despite the clear and almost complete assignment previously achieved (35). These resonances, which correspond to residues equally distributed along the josephin domain, were genuinely not observed in the full-length ataxin-3 spectrum and undetectable in the residue specific selectively labelled samples suggesting that the full-length protein has different exchange regimes from the isolated domain.

Our assignment gives us now, for the first time, an overall picture of the structure of this protein: despite being overall highly flexible, the C-terminal tail of ataxin-3 contains several stretches of well-defined secondary structure. The three UIMs are helical in agreement with a previous study of a construct containing UIM1 and 2 (11). In addition to these regions, a helical tract was observed also between residues 278 and 289 which directly precede the polyQ tract. The identification of a stable α-helix in this region was recently suggested by a crystallographic study (36) on a construct containing ataxin-3 residues 278-305 fused to MBP and a short C-terminal crystallization tag. Interestingly, two different crystals were obtained in this study. One included the monomeric MBP with the ataxin-3 region 278-305 exposed to the solvent. In this structure, residues 278-284 form two helical turns, while the other residues could not be modelled because of absence of electron density. In the second crystal, the protein forms a dimer in which the tract 278-304 is helical, partially shielded from the solvent and stabilised by contacts with MBP. We performed our structural analysis of full-length ataxin-3 in solution, under near-physiologic conditions and without using any tag that could potentially interfere with the structure of the protein. We can thus conclusively affirm that the 278-289 tract forms a helical conformation in solution when in a monomeric state. It was not possible to obtain sequential assignment and hence estimate the structure of the N-terminal residues of the polyQ repeats (292-297). It is likely that these residues are more rigid than the other repeats, as also suggested by the fact that the resonances of these residues are below the detection threshold of 5D experiments. The possibility that they are in an intermediate exchange regime is ruled out by the fact that the associated H, N, C_α_ and C_β_ resonances were detectable in the HNCACB experiment, albeit not sequentially assignable because of severe overlap of their C_α_ and C_β_ resonances with those of other residues. However, since minor helical tendency was observed within the following polyQ repeats this could suggest that this region has a tendency to adopt a helical conformation stabilized by the preceding residues. This hypothesis would be supported by the crystal structure of exon I of huntingtin (37), another well characterised member of the polyQ family, where the whole region is helical.

## Conclusions

In conclusion, the ability to study the structural behaviour of this protein paves the way to the investigation of interactions with other proteins. An obvious candidate is the interaction with the valosin-containing protein which was mapped in the basic motif RKRR in the C-terminal tail (38). Our work will also potentially be important for the advance of ataxin-3 aggregation since a number of studies have shown that the regions immediately N-and C-terminal to the polyQ repeats play an important role in modulating protein self-assembly (39-44).

## Acknowledgements

We are indebted with the MRC NMR facilities at The Francis Crick Institute London. CIISB research infrastructure project LM2015043 funded by MEYS CR is also gratefully acknowledged.

## References

1 Yoshimura, Y., N. V. Kulminskaya, and F. A. Mulder. 2015. Easy and unambiguous sequential assignments of intrinsically disordered proteins by correlating the backbone 15N or 13C’ chemical shifts of multiple contiguous residues in highly resolved 3D spectra. J Biomol NMR 61:109–121.

2 Isaksson, L., M. Mayzel, M. Saline, A. Pedersen, J. Rosenlow, B. Brutscher, B. G. Karlsson, and V. Y. Orekhov. 2013. Highly efficient NMR assignment of intrinsically disordered proteins: application to B- and T cell receptor domains. PLoS One 8:e62947.

3 Goradia, N., C. Wiedemann, C. Herbst, M. Gorlach, S. H. Heinemann, O. Ohlenschlager, and R. Ramachandran. 2015. An approach to NMR assignment of intrinsically disordered proteins. Chemphyschem 16:739–746.

4 Paulson, H. L., M. K. Perez, Y. Trottier, J. Q. Trojanowski, S. H. Subramony, S. S. Das, P. Vig, J. L. Mandel, K. H. Fischbeck, and R. N. Pittman. 1997. Intranuclear inclusions of expanded polyglutamine protein in spinocerebellar ataxia type 3. Neuron 19:333–344.

5 Masino, L., V. Musi, R. P. Menon, P. Fusi, G. Kelly, T. A. Frenkiel, Y. Trottier, and A. Pastore. 2003. Domain architecture of the polyglutamine protein ataxin-3: a globular domain followed by a flexible tail. FEBS Lett 549:21–25.

6 Ellisdon, A. M., B. Thomas, and S. P. Bottomley. 2006. The two-stage pathway of ataxin-3 fibrillogenesis involves a polyglutamine-independent step. J Biol Chem 281:16888–16896.

7 Saunders, H. M., D. Gilis, M. Rooman, Y. Dehouck, A. L. Robertson, and S. P. Bottomley. 2011. Flanking domain stability modulates the aggregation kinetics of a polyglutamine disease protein. Protein Sci 20:1675–1681.

8 Doss-Pepe, E. W., E. S. Stenroos, W. G. Johnson, and K. Madura. 2003. Ataxin-3 interactions with rad23 and valosin-containing protein and its associations with ubiquitin chains and the proteasome are consistent with a role in ubiquitin-mediated proteolysis. Mol Cell Biol 23:6469–6483.

9 Scaglione, K. M., E. Zavodszky, S. V. Todi, S. Patury, P. Xu, E. Rodriguez-Lebron, S. Fischer, J. Konen, A. Djarmati, J. Peng, J. E. Gestwicki, and H. L. Paulson. 2011. Ube2w and ataxin-3 coordinately regulate the ubiquitin ligase CHIP. Mol Cell 43:599–612.

10 Nicastro, G., L. Masino, V. Esposito, R. P. Menon, A. De Simone, F. Fraternali, and A. Pastore. 2009. Josephin domain of ataxin-3 contains two distinct ubiquitin-binding sites. Biopolymers 91:1203–1214.

11 Song, A. X., C. J. Zhou, Y. Peng, X. C. Gao, Z. R. Zhou, Q. S. Fu, J. Hong, D. H. Lin, and H. Y. Hu. 2010. Structural transformation of the tandem ubiquitin-interacting motifs in ataxin-3 and their cooperative interactions with ubiquitin chains. PLoS One 5:e13202.

12 Delaglio, F., S. Grzesiek, G. W. Vuister, G. Zhu, J. Pfeifer, and A. Bax. 1995. NMRPipe: a multidimensional spectral processing system based on UNIX pipes. J Biomol NMR 6:277–293.

13 Kazimierczuk, K., W. Kozminski, and I. Zhukov. 2006. Two-dimensional Fourier transform of arbitrarily sampled NMR data sets. J Magn Reson 179:323–328.

14 Kazimierczuk, K., A. Zawadzka, W. Kozminski, and I. Zhukov. 2006. Random sampling of evolution time space and Fourier transform processing. J Biomol NMR 36:157–168.

15 Kazimierczuk, K., A. Zawadzka, and W. Kozminski. 2009. Narrow peaks and high dimensionalities: exploiting the advantages of random sampling. J Magn Reson 197:219–228.

16 Vranken, W. F., W. Boucher, T. J. Stevens, R. H. Fogh, A. Pajon, M. Llinas, E. L. Ulrich, J. L. Markley, J. Ionides, and E. D. Laue. 2005. The CCPN data model for NMR spectroscopy: development of a software pipeline. Proteins 59:687–696.

17 Helmus, J. J., and C. P. Jaroniec. 2013. Nmrglue: an open source Python package for the analysis of multidimensional NMR data. J Biomol NMR 55:355–367.

18 Masino, L., G. Nicastro, A. De Simone, L. Calder, J. Molloy, and A. Pastore. 2011. The Josephin domain determines the morphological and mechanical properties of ataxin-3 fibrils. Biophys J 100:2033–2042.

19 Lescop, E., T. Kern, and B. Brutscher. 2010. Guidelines for the use of band-selective radiofrequency pulses in hetero-nuclear NMR: example of longitudinal-relaxation-enhanced BEST-type 1H-15N correlation experiments. J Magn Reson 203:190–198.

20 Motackova, V., J. Novacek, A. Zawadzka-Kazimierczuk, K. Kazimierczuk, L. Zidek, H. Sanderova, L. Krasny, W. Kozminski, and V. Sklenar. 2010. Strategy for complete NMR assignment of disordered proteins with highly repetitive sequences based on resolution-enhanced 5D experiments. J Biomol NMR 48:169–177.

21 MacRaild, C. A., M. Zachrdla, D. Andrew, B. Krishnarjuna, J. Novacek, L. Zidek, V. Sklenar, J. S. Richards, J. G. Beeson, R. F. Anders, and R. S. Norton. 2015. Conformational dynamics and antigenicity in the disordered malaria antigen merozoite surface protein 2. PLoS One 10:e0119899.

22 Nyarko, A., Y. Song, J. Novacek, L. Zidek, and E. Barbar. 2013. Multiple recognition motifs in nucleoporin Nup159 provide a stable and rigid Nup159-Dyn2 assembly. J Biol Chem 288:2614–2622.

23 Orban-Nemeth, Z., M. A. Henen, L. Geist, S. Zerko, S. Saxena, J. Stanek, W. Kozminski, F. Propst, and R. Konrat. 2014. Backbone and partial side chain assignment of the microtubule binding domain of the MAP1B light chain. Biomol NMR Assign 8:123–127.

24 Solyom, Z., M. Schwarten, L. Geist, R. Konrat, D. Willbold, and B. Brutscher. 2013. BEST-TROSY experiments for time-efficient sequential resonance assignment of large disordered proteins. J Biomol NMR 55:311–321.

25 Marsh, J. A., V. K. Singh, Z. Jia, and J. D. Forman-Kay. 2006. Sensitivity of secondary structure propensities to sequence differences between alpha- and gamma-synuclein: implications for fibrillation. Protein Sci 15:2795–2804.

26 Li, P., K. E. Huey-Tubman, T. Gao, X. Li, A. P. West, Jr., M. J. Bennett, and P. J. Bjorkman. 2007. The structure of a polyQ-anti-polyQ complex reveals binding according to a linear lattice model. Nat Struct Mol Biol 14:381–387.

27 Masino, L., G. Kelly, K. Leonard, Y. Trottier, and A. Pastore. 2002. Solution structure of polyglutamine tracts in GST-polyglutamine fusion proteins. FEBS Lett 513:267–272.

28 Nicastro, G., R. P. Menon, L. Masino, P. P. Knowles, N. Q. McDonald, and A. Pastore. 2005. The solution structure of the Josephin domain of ataxin-3: structural determinants for molecular recognition. Proc Natl Acad Sci U S A 102:10493–10498.

29 Maciejewski, M. W., D. Liu, R. Prasad, S. H. Wilson, and G. P. Mullen. 2000. Backbone dynamics and refined solution structure of the N-terminal domain of DNA polymerase beta. Correlation with DNA binding and dRP lyase activity. J Mol Biol 296:229–253.

30 Burnett, B., F. Li, and R. N. Pittman. 2003. The polyglutamine neurodegenerative protein ataxin-3 binds polyubiquitylated proteins and has ubiquitin protease activity. Hum Mol Genet 12:3195–3205.

31 Chai, Y., S. S. Berke, R. E. Cohen, and H. L. Paulson. 2004. Poly-ubiquitin binding by the polyglutamine disease protein ataxin-3 links its normal function to protein surveillance pathways. J Biol Chem 279:3605–3611.

32 Chow, M. K., J. P. Mackay, J. C. Whisstock, M. J. Scanlon, and S. P. Bottomley. 2004. Structural and functional analysis of the Josephin domain of the polyglutamine protein ataxin-3. Biochem Biophys Res Commun 322:387–394.

33 Chow, M. K., H. L. Paulson, and S. P. Bottomley. 2004. Destabilization of a non-pathological variant of ataxin-3 results in fibrillogenesis via a partially folded intermediate: a model for misfolding in polyglutamine disease. J Mol Biol 335:333–341.

34 Nicastro, G., M. Habeck, L. Masino, D. I. Svergun, and A. Pastore. 2006. Structure validation of the Josephin domain of ataxin-3: conclusive evidence for an open conformation. J Biomol NMR 36:267–277.

35 Nicastro, G., L. Masino, T. A. Frenkiel, G. Kelly, J. McCormick, R. P. Menon, and A. Pastore. 2004. Assignment of the 1H, 13C, and 15N resonances of the Josephin domain of human ataxin-3. J Biomol NMR 30:457–458.

36 Zhemkov, V. A., A. A. Kulminskaya, I. B. Bezprozvanny, and M. Kim. 2016. The 2.2-Angstrom resolution crystal structure of the carboxy-terminal region of ataxin-3. FEBS Open Bio 6:168– 178.

37 Kim, M. W., Y. Chelliah, S. W. Kim, Z. Otwinowski, and I. Bezprozvanny. 2009. Secondary structure of Huntingtin amino-terminal region. Structure 17:1205–1212.

38 Rao, M. V., D. R. Williams, S. Cocklin, and P. J. Loll. 2017. Interaction between the AAA(+) ATPase p97 and its cofactor ataxin3 in health and disease: Nucleotide-induced conformational changes regulate cofactor binding. J Biol Chem 292:18392–18407.

39 Bhattacharyya, A., A. K. Thakur, V. M. Chellgren, G. Thiagarajan, A. D. Williams, B. W. Chellgren, T. P. Creamer, and R. Wetzel. 2006. Oligoproline effects on polyglutamine conformation and aggregation. J Mol Biol 355:524–535.

40 Darnell, G., J. P. Orgel, R. Pahl, and S. C. Meredith. 2007. Flanking polyproline sequences inhibit beta-sheet structure in polyglutamine segments by inducing PPII-like helix structure. J Mol Biol 374:688–704.

41 Thakur, A. K., M. Jayaraman, R. Mishra, M. Thakur, V. M. Chellgren, I. J. Byeon, D. H. Anjum, R. Kodali, T. P. Creamer, J. F. Conway, A. M. Gronenborn, and R. Wetzel. 2009. Polyglutamine disruption of the huntingtin exon 1 N terminus triggers a complex aggregation mechanism. Nat Struct Mol Biol 16:380–389.

42 Eftekharzadeh, B., A. Piai, G. Chiesa, D. Mungianu, J. Garcia, R. Pierattelli, I. C. Felli, and X. Salvatella. 2016. Sequence Context Influences the Structure and Aggregation Behavior of a PolyQ Tract. Biophys J 110:2361–2366.

43 Shen, K., B. Calamini, J. A. Fauerbach, B. Ma, S. H. Shahmoradian, I. L. Serrano Lachapel, W. Chiu, D. C. Lo, and J. Frydman. 2016. Control of the structural landscape and neuronal proteotoxicity of mutant Huntingtin by domains flanking the polyQ tract. Elife 5.

44 Totzeck, F., M. A. Andrade-Navarro, and P. Mier. 2017. The Protein Structure Context of PolyQ Regions. PLoS One 12:e0170801.

